# Hematopoietic p53 loss cell-extrinsically defines an immune infiltrated microenvironment in leukemia and pre-leukemia

**DOI:** 10.1101/2020.03.22.002774

**Authors:** Ryan R. Posey, Jonathan D. Lee, Sean Clohessy, Lourdes M. Mendez, Pier Paolo Pandolfi

**Affiliations:** Cancer Research Institute, Beth Israel Deaconess Cancer Center, Department of Medicine and Pathology, Beth Israel Deaconess Medical Center, Harvard Medical School, Boston, MA, USA; Ludwig Center at Harvard, Harvard Medical School, Boston, MA, USA; Department of Molecular Biotechnology and Health Sciences, Molecular Biotechnology Center, University of Turin, Italy

## Abstract

*TP53* is the most frequently mutated gene in human cancers. In Acute Myeloid Leukemia (AML) and Clonal Hematopoiesis of Indeterminate Potential (CHIP), it is one of several recurrent genetic alterations. Despite multiple recent therapeutic advances for AML, *TP53* mutated AML is associated with resistance to currently approved therapies and thus, a very poor prognosis. Emerging evidence suggests that mutations in *TP53* may be a predictor of positive response to immunotherapy. To model cell - extrinsic consequences of hematopoietic p53 loss, we generated bone marrow chimeric mice bearing *p53*^-/-^ and congenic wild type cells. Following reconstitution, we observed increased levels of wild type CD8+ and CD4+ T cells in mice transplanted with *p53*^-/-^ hematopoietic cells compared to controls. In addition, we observed a change in the frequency of T cell subsets in *p53*^-/-^ chimeras including an increase in Tregs. To determine if these alterations were mirrored in the leukemic setting, we next generated *p53*^-/-^;*nRas^G12D^* leukemia. While the bone marrow of *p53*^-/-^;*nRas^G12D^* leukemia showed the presence of both T and B lymphocytes, MLL-AF9 showed a near complete absence of lymphocytes, akin to ‘immune-infiltrated’ and ‘immune-desert’ phenotypes seen in solid tumors. These data clearly demonstrate a causal cell-extrinsic effect of hematopoietic p53 loss on the immune system, both in the context of leukemia and preleukemic states. Modeling AML genetics in murine models serves as a powerful tool to define the association between genetic drivers and immune subtypes of AML towards precise patient stratification critical for the application of emerging targeted and immune therapies.

**Statement of Significance:** *TP53* mutations are frequent in both AML and CHIP, and are associated with both resistance to therapy as well as very poor prognosis. We provide evidence to investigate the immunotherapy as a treatment option for this subgroup of AML.

## Introduction

*TP53* is a frequently mutated gene in both Acute Myeloid Leukemia (AML) and Clonal Hematopoiesis of Indeterminate Potential (CHIP)[1–4]. According to the prognostic classification system described by Döhner et al., *TP53* mutations in AML are associated with very poor outcome as well as complex karyotype[5]. This poor outcome is believed to be a consequence of the poor responsiveness to standard therapy regimens, such as 7+3 induction, treatment with hypomethylating agents, and allogeneic bone marrow transplant (BMT)[6]. Similarly, when observed in CHIP, patients with *TP53* mutations have the highest risk of progression to AML, with 100% developing AML in one cohort[7]. Unsurprisingly, the presence of *TP53* mutations several years prior to the diagnosis is associated with the development therapy-related AML (t-AML), following cytotoxic chemotherapy or radiotherapy[8, 9].

CHIP has recently been identified as a risk factor for cardiovascular disease [10]. Utilizing competitive bone marrow transplantation in mice as a model of CHIP, multiple studies have shown that TET2 deficiency accelerates atherosclerosis as a consequence of aberrant IL-1ß-dependent pro-inflammatory signaling in macrophages[10–12]. Interestingly, using a similar model, DNMT3A mutations were found to have an altered Th17:Treg ratio in patients, suggesting a similar pro-inflammatory phenotype mediated by an altogether different mechanism[13].

Given the frequency of *TP53* mutations in CHIP, we hypothesized that mutations in this tumor suppressor could similarly alter the surrounding immune landscape. In line with this, recent *in silico* efforts have suggested *TP53*-mutated AML is strongly associated with an immune-infiltrated signature in both the BeatAML and TCGA cohorts[14–17]. Modeling *TP53* CHIP disorders and AML progressing from CHIP as previously described in the literature, we describe cell-extrinsic expansion of lymphocytes as a consequence of hematopoietic *Trp53* loss and a resulting ‘immune infiltrated’ phenotype in a *Trp53*^-/-^;*Nras^G12D^* murine model of AML[18].

## Results

### Hematopoietic p53 loss drives subset-specific expansion of T lymphocytes

To model hematopoietic p53 loss, we generated mixed bone marrow chimeras of either *Trp53*^-/-^ or wildtype cells (CD45.2) and congenic wild-type cells (CD45.1) into lethally irradiated, congenic, wild-type recipients (CD45.1) (Figure 1A). Peripheral blood T cells were tracked in the peripheral blood over time by regular bleedings until euthanasia at 10 weeks. During this time, these mice display no symptoms of disease and appeared otherwise healthy. Following reconstitution, we observed an increase in the number of total CD45.1+, but not CD45.2+ T cells (Figure 1B). Consistent with previous reports, *Trp53*^-/-^ leukocytes outcompete wild-type cells over time (Figure 1C). This effect was profound in the peripheral blood of *Trp53*^-/-^ chimeras and increased over time to approximately double the frequency of T cells (Figure 1D). A positive correlation was observed between outcompetition by *Trp53*^-/-^ cells and the increase in wild-type T cells (Supplementary Figure 1). At time of euthanasia, the spleens of these mice showed a similar increase in the numbers of T cells, suggesting a systemic effect driven by *Trp53* loss (Figure 1B). Interestingly, we did not observe increased frequencies of either CD4+ or CD8+ central memory or CD8+ effector/effector memory *Trp53^+/+^* T cells; rather, both CD4+ and CD8+ naïve, CD4+ effector memory, and CD4+ Treg subsets were expanded (Figure 2A). These data suggest primarily a naïve, rather than anergic T cell phenotype, implicating other possible mechanisms of dysfunction aside from overexpression of checkpoint molecules. Given the proportional expansion of CD4+ Tregs, it is further possible that effector function is restrained by these cells, limiting alloreactivity. Additionally, in the short-term following reconstitution, *Trp53*^-/-^ chimeras displayed a decreased proportion of CD8+ effector/effector memory cells, further suggesting abrogated effector T cell differentiation (Figure 2E). Several weeks later, an increased proportion of Tregs was observed, although this effect was only observed at a single time point (Figure 2F). These effects were consistent when a small number of cKit+, Sca-1+, Lineage-cells were transplanted into sub-lethally irradiated recipients (Supplementary Figure 2A). However, we did not observe decreased effector/effector memory T cells shortly after reconstitution with this alternative method of transplantation (Supplementary Figure 2B). Interestingly, we observed increased levels of the T cell stemness marker Sca-1 at 8- and 10-weeks posttransplantation in the peripheral blood of recipients of *Trp53*^-/-^ KSL, but not at 6 weeks (Supplementary Figure 3A-B). This increase was concurrent with the expansion of all T cells, suggesting aberrant T cell stemness as a key driver of this effect. In addition, we observed an increase in the ratio of granulocytes to monocytes in both the peripheral blood and spleens of mixed bone marrow chimeras (Figure 2B,D) and a significant decrease in the frequency of *Trp53^+/+^* monocytes within the spleen. This observation suggests aberrant pro-inflammatory signaling as a mechanism of these observed changes, consistent with a role of p53 in modulating systemic inflammation and neutrophil homeostasis[19]. No statistically significant changes were observed within p53-null cells (CD45.2+, CD45.1-) and their wild-type counterparts in levels of total CD3+ or CD4+ T cells; however, a modest but significant increase in the number of CD8+ T cells was observed (Figure 3A,C). Additionally, a cell-intrinsic bias was observed with both CD4+ and CD8+ effector/effector memory and central memory T cell subsets expanded (Figure 3A).

**Figure 1:**
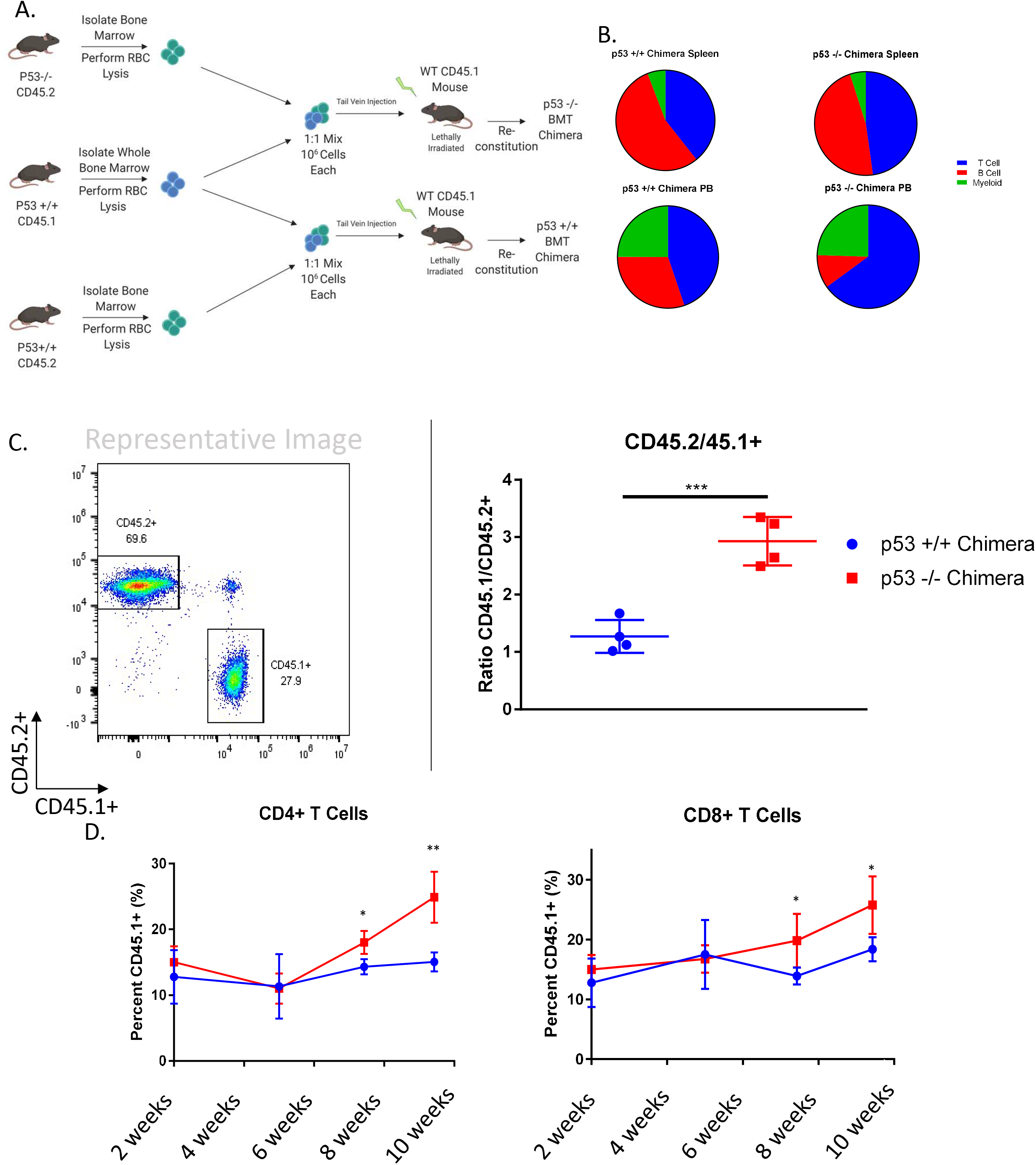
p53 Loss Cell-Extrinsically Expands T Cells. (A) Schematic showing generation of mixed bone marrow chimeras. (B) Pie charts representing changes in overall composition of spleen and periphery as a consequence of *Trp53*^-/-^ bone marrow transplantation. (C) Representative flow plots and outcompetition of *Trp53*^-/-^ cells (CD45.2+) as compared to congenic Bl6 controls. (D) Increase in CD45.1+ T cells over 10 weeks.

**Figure 2:**
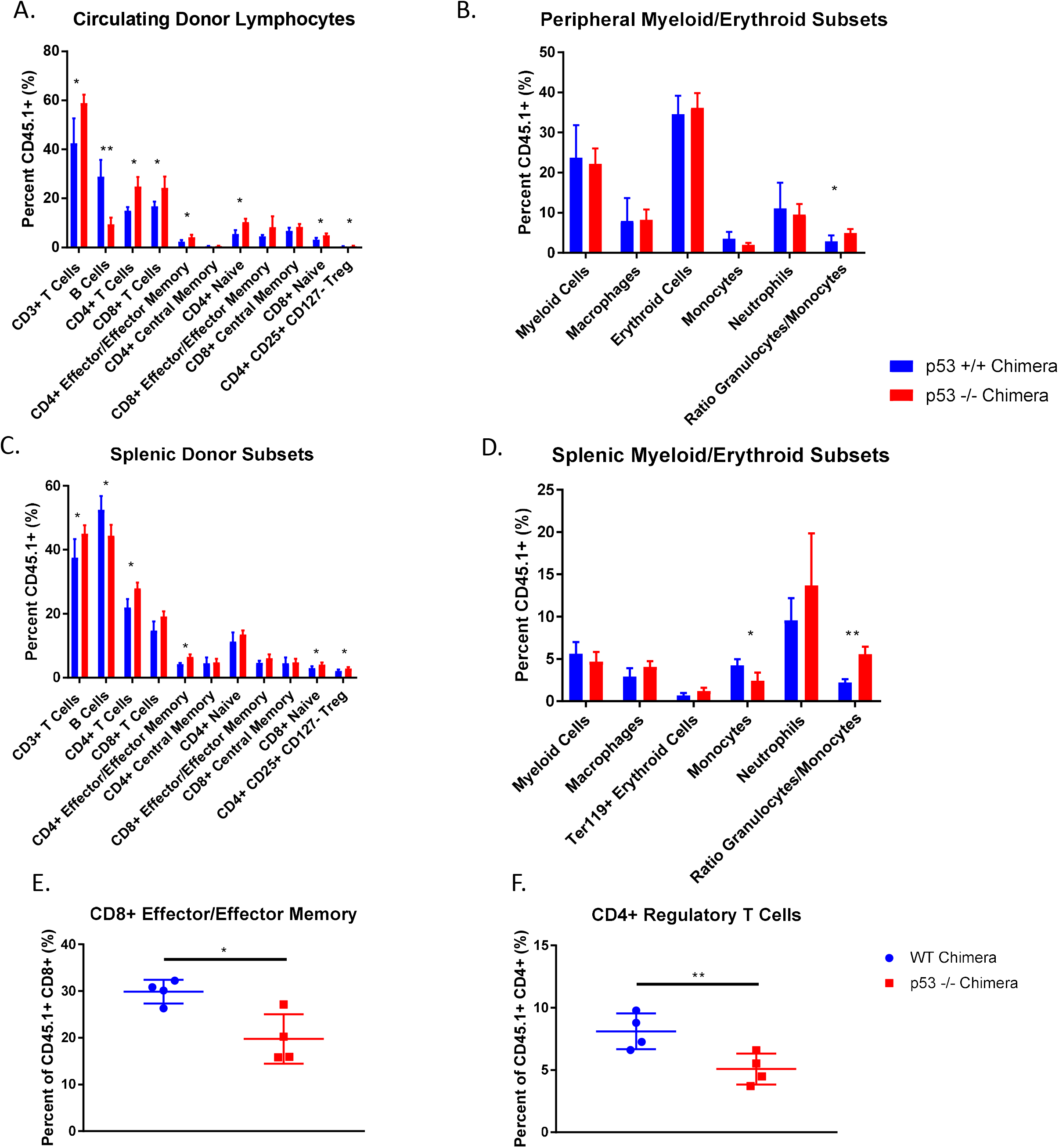
Cell-Extrinsic subset-specific changes as a consequence of *Trp53* loss. Peripheral Blood (A) and Spleen (C) T cell subsets as compared with Bl6 control chimeras. Peripheral Blood (B) and Spleen (D) Myeloid/Erythroid subsets within *Trp53*^-/-^ chimeras as compared to Bl6 controls. (E) CD8+ Effector/Effector memory T cells as a percent of total CD8+ T cells at 3 weeks posttransplantation. (F) CD4+ Regulatory T cells as a percentage of total CD4+ T cells at 5 weeks posttransplantation.

**Figure 3:**
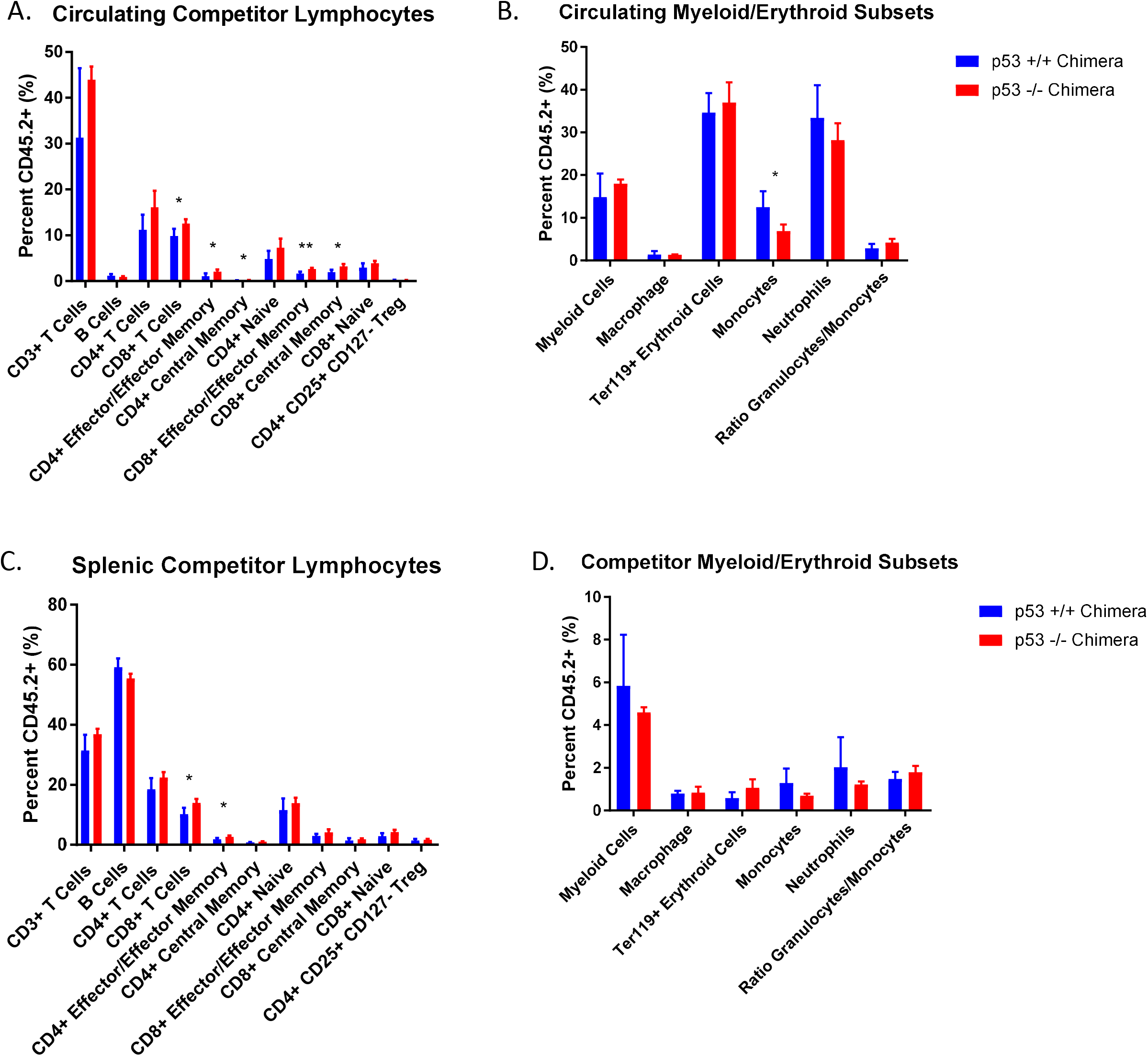
Cell-Intrinsic Changes as a consequence of *Trp53* loss. (A) Circulating *Trp53*^-/-^ T cell and myeloid (B) subsets as a fraction of total CD45.2+ T leukocytes. (C) Splenic T and myeloid subsets (D) as a fraction of total CD45.2+ leukocytes.

No other significant differences were observed in the frequencies of CD4+ and CD8+ T cells. In addition, a significant decrease was observed in the frequencies of circulating, but not splenic monocytes, although this change was not reflected in the ratio of granulocytes to monocytes (Figure 3B,D). Taken together, these data strongly support a cell-extrinsic effect of *Trp53* loss in expanding the global pool of T lymphocytes.

### Identification of Immune Infiltrated and Immune Desert Subsets of AML

Next, we sought to determine if this effect was relevant to the leukemia setting. We therefore generated both p53-independent *MLL-AF9* and p53-dependent *Trp53*^-/-^;*Nras^G12D^* models of AML. MLL-AF9 leukemia was generated by retrovirus-mediated overexpression using the MSCV-MLL-AF9-IRES-GFP vector (MIG-MF9). The *Trp53*^-/-^;*Nras^G12D^* model was generated by retrovirus-mediated overexpression of Nras using the pBabe-MSCV-Nras-RFP. The median survival time for mice transplanted with MLL-AF9 leukemia was 30 days and 38 days for *Trp53*^-/-^;*Nras^G12D^*. In line with published reports, MLL-AF9 immunophenotypically resembled a monocytic leukemia, and expressed the markers CD11b and Ly6C but not Ly6G (Figure 4B). In contrast, *Trp53*^-/-^;*Nras^G12D^* leukemia resembled an erythroleukemia, with ~20% of blasts expressing the erythroid marker Ter119 (Figure 4B, Supplementary Figure 4A). However, the majority of the leukemia was negative for any of the lineage antigens tested. Concordant with the literature, both diseases were serially transplantable. Transplantation of small numbers of RFP+ megakaryocyte-erythroid progenitors into sub-lethally irradiated recipients engrafted in all recipients, consistent with the role of these progenitors as leukemia initiating cells (data not shown). Flow cytometry (Supplementary Figure 4B) and hematoxylin and eosin staining of spleen and liver (Supplementary Figure 4C) showed extensive hepatosplenomegaly, in line with previously published reports.

**Figure 4:**
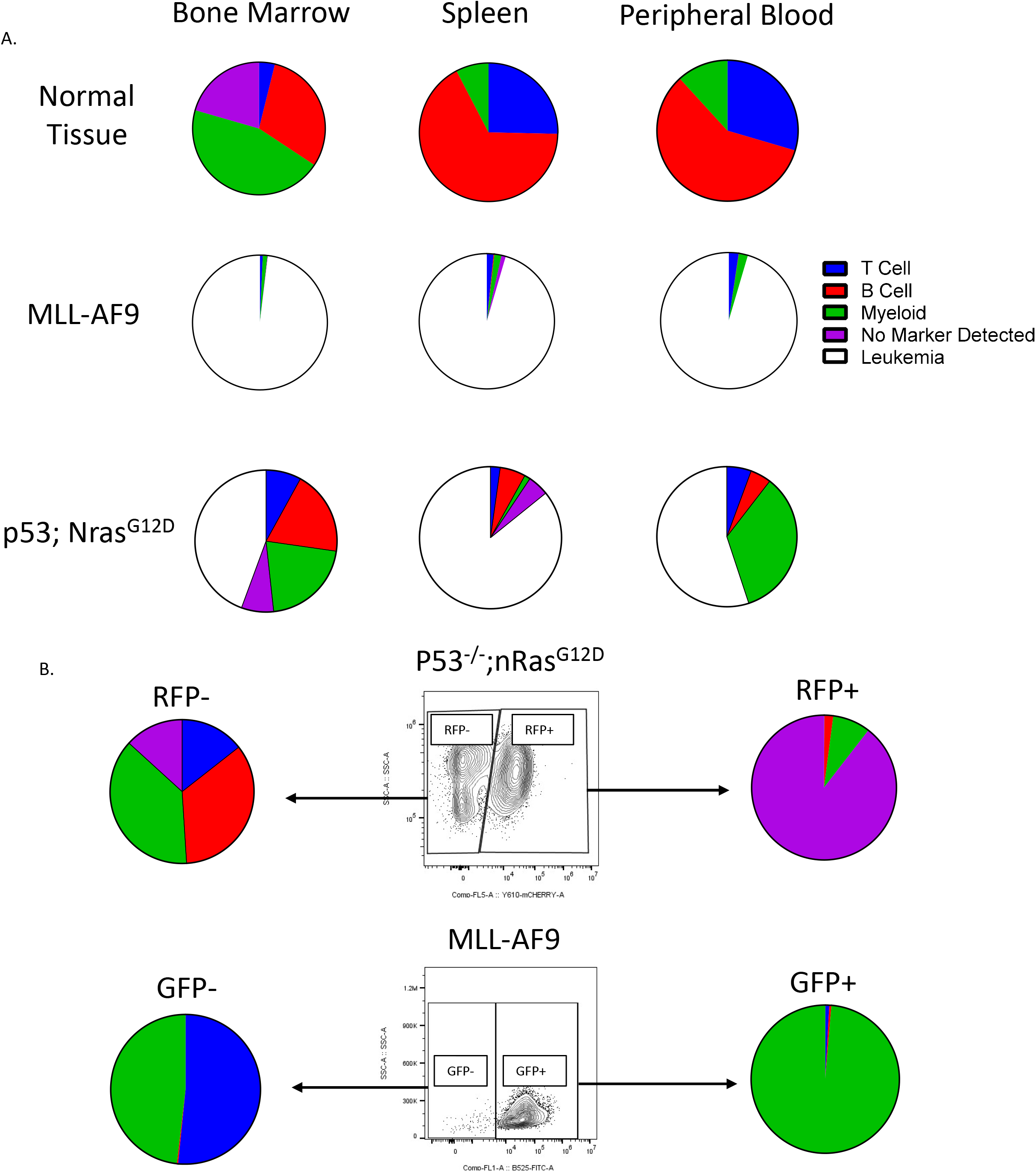
Identification of ‘Immune-Infiltrated’ and ‘Immune Desert’ tumors in murine models of AML. (A) Immune landscape across several hematopoietic tissues in normal tissue and MLL-AF9 or *Trp53*^-/-^;*Nras^G12D^* leukemia. (B) Detailed immunophenotyping of *Trp53*^-/-^;*Nras^G12D^* and MLL-AF9 leukemia.

To account for differences in burden of disease and altered kinetics within the various hematopoietic niches infiltrated by either subtype of AML, we examined primary recipient mice moribund at time of euthanasia. MLL-AF9 recipients displayed almost a complete global absence of normal immune cells, as observed in the blood, spleen, and bone marrow (Figure 4A). This phenotype was reminiscent of an ‘immune desert’ seen in solid tumors and is in accordance with previous reports. This finding is unsurprising, as immune deserts in solid tumors are predictors of poor response to immunotherapy, and the majority of AMLs do not respond to checkpoint inhibition as a monotherapy[20, 21]. In contrast, *Trp53*^-/-^;*Nras^G12D^* leukemia showed the presence of infiltrating immune cells within the blood, spleen, and bone marrow at time of euthanasia (Figure 4A). This was reminiscent of an ‘immune-infiltrated’ phenotype observed in solid tumors, suggesting that this subtype of AML may be susceptible to immunotherapy as is the case with solid tumors. We therefore establish that such phenotypes are present in AML.

## Discussion

*TP53* is a recurrently mutated gene in AML and CHIP, and a predictor of very poor prognosis. While the immune cell repertoire of solid tumors has been extensively described in both patients and murine models, no such efforts have yet been made in AML. Consequently, we describe immune infiltrated and immune desert subsets of AML as a direct consequence of their genetic aberrations. This is in accordance with previous *in silica* analyses implicating *TP53*-mutated AML as an immune infiltrated tumor. Further, owing to the ability to model single-gene perturbations in the hematopoietic system using bone marrow transplantation, the effect of individual driver mutations on surrounding immune repertoire can be described. By two independent transplantation methods, we observed cell-extrinsic increases in the frequencies of circulating, *Trp53^+/+^* T lymphocytes, as well as an increase in the ratio of monocytes to granulocytes. This suggests, as seen in solid tumors, that loss of this tumor suppressor can cause pro-inflammatory and leukemia-supporting inflammation, in line with this description of *TP53*-mutated AML. In addition, previous studies examining patients have directly observed increased levels of PD-1 expression within this subtype [22–24], further suggesting *TP53*-mutated AML may be susceptible to immunotherapy. Taken together, these data argue for further studies investigating the possibility of treating *TP53*-mutant AML with various forms of immunotherapy.

## Methods

### Animal Husbandry

The Institutional Animal Care and Use Committee (IACUC) of Beth Israel Deaconess Medical Center granted approval for all mouse experiments described here. The transgenic mice used in this study were obtained from the Jackson Laboratories (*B6.129S2-Trp53tm1Tyj/J*, stock #002101; *B6.SJL-Ptprca Pepcb /BoyJ*, stock #002014).

### Bone Marrow Transplantation

Recipient CD45.1+ mice were lethally irradiated with one dose of γ-irradiation (950 cGy). Donor CD45.2+ bone marrow was obtained from either *Trp53^+/+^* and *Trp53*^-/-^ mice at 10 weeks of age and one CD45.1+ mouse of the same age. Recipients were transplanted with 2×10^6^ whole bone marrow cells at a 1:1 ratio four hours following irradiation by tail vein injection (TVI). Alternatively, recipients were sublethally irradiated with one dose of γ-irradiation (650 cGy) and transplanted with cultured KSL (TVI).

### Murine Leukemia Models

The MLL-AF9 model was generated as previously described[25]. Whole bone marrow nucleated cells were isolated by crushing tibias, femurs, pelves, and spines and passing through a 100 μm filter. Cells were spun down and lysed briefly using ACK lysis buffer (ThermoFisher catalog #A1049201) before being diluted 1:10 in a solution of 2% FBS. These cells were stained with a commercial lineage cocktail (Biolegend catalog #133310), anti-Sca-1-PE (Biolegend catalog #108108), and anti-cKit-APC (Biolegend catalog #313206). Sorted cells were cultured in StemSpan (StemCell catalog #09650) with 10 ng/μL Il-3 and Il-6 and 50 ng/μL SCF, TPO, and Flt3L.

### Plasmids

MSCV-MLL-AF9-IRES-GFP was a kind gift from David Sykes. The pBabe-MSCV-Nras^G12D^-Ires-RFP vector was cloned by excision from a donor Nras^12D^ open reading frame plasmid (Addgene # 14725) using BamHI. A base pMSCV-pBABEMCS-IRES-RFP (Addgene #33337) was digested using BamHI and ligated to the excised product. The resulting plasmid was then sequence validated.

### Murine Retrovirus Infection and Transplantation

Recombinant murine retroviruses were produced as previously described[25]. Briefly, 293T cells were transfected using Lipofectamine 2000 (ThermoFisher catalog # 11668019), pCL-Eco, and one of the following transfer plasmids: pBabe-MSCV-RFP, pBabe-MSCV-Nras^G12D^, MSCV-MLL-AF9-IRES-GFP. Supernatant was changed one day following transfection, collected at 48 hours post-infection, and concentrated using a 100K centrifuge column (Millipore-Sigma catalog # UFC910008). Cells were spin-infected with 200 μL of concentrated retrovirus and 8 μg/mL polybrene. Following infection, cells were washed twice with PBS before being counted and injected into recipients.

### Analysis of Peripheral Blood and Tissue Samples

Blood was collected from mice by submandibular bleeding into EDTA collection tubes. This EDTA-anticoagulated whole blood was lysed using ACK lysis buffer for 5 minutes, then diluted 1:10 in a solution of 2% FBS, centrifuged, and lysed again for another 5 minutes. Cells were then resuspended in 2% FBS and stained for cell populations as described below.

### Flow Cytometry

Cellular subpopulations were analyzed by flow cytometry using the following antibodies: PE-conjugated anti-CD3, FITC-conjugated anti-CD19, PerCP-Cy5.5-conjugated anti-CD45.1, Pacific Blue-conjugated anti-CD45.2, APC-Cy7-conjugated anti-B220, PE-Cy7-conjugated anti-CD11b, FITC-conjugated anti-CD4, PE-conjugated anti-CD8, APC-Cy7-conjugated anti-CD44, BV421-conjugated anti-CD127, APC-conjugated anti-CD25, BV605-conjugated anti-CD62L, PE-Cy7-conjugated anti-CD45.2, BV510-conjugated anti-CD45.2, PE-conjugated anti-CD4, PE-Cy7-conjugated anti-CD11b, PE-conjugated anti-Ly6C, APC-conjugated anti-Ly6G, BV605-conjugated anti-CD11c, APC-Cy7-conjugated anti-F4/80, PE-Cy7-conjugated anti-CD206, FITC-conjugated anti-Ly6C, PE-conjugated anti-CD11b, APC-conjugated anti-Ter119, PE-conjugated anti-Sca-1, APC-conjugated anti-CD117, Pacific Blue-conjugated lineage cocktail. HPSCs were identified as Lineage-, cKit+, Sca1+; monocytes, as CD45+, CD11b+, Ly6C^hi^, Ly6G+; neutrophils, as CD45+, CD11b+, Ly6C^int^, Ly6g+, T lymphocytes as CD45+, CD11b-, B220-, CD3+; B lymphocytes as CD45+, CD11b-, CD3-,CD19+, B220+; macrophages as CD45+, CD11B+, F4/80^Hi^; M1-like macrophages as CD45+, CD11b+, F4/80^Hi^, CD11c^hi^, CD206^lo^, M2-like macrophages as CD45+, CD11b+, F4/80^Hi^, CD11c^lo^, CD206^hi^, effector memory T cells as CD44^Hi^ CD62L-, naïve T cells as CD44^Lo^ CD62L+, and regulatory T cells as CD4+ CD127Lo CD25+. A CytoFLEX LX flow cytometer (Beckman Coulter) was used for data acquisition. Data were analyzed with FlowJo Software. For sorting, a FACSAria II (BD Biosciences) was used.

---

**Supplemental Figure 1:**
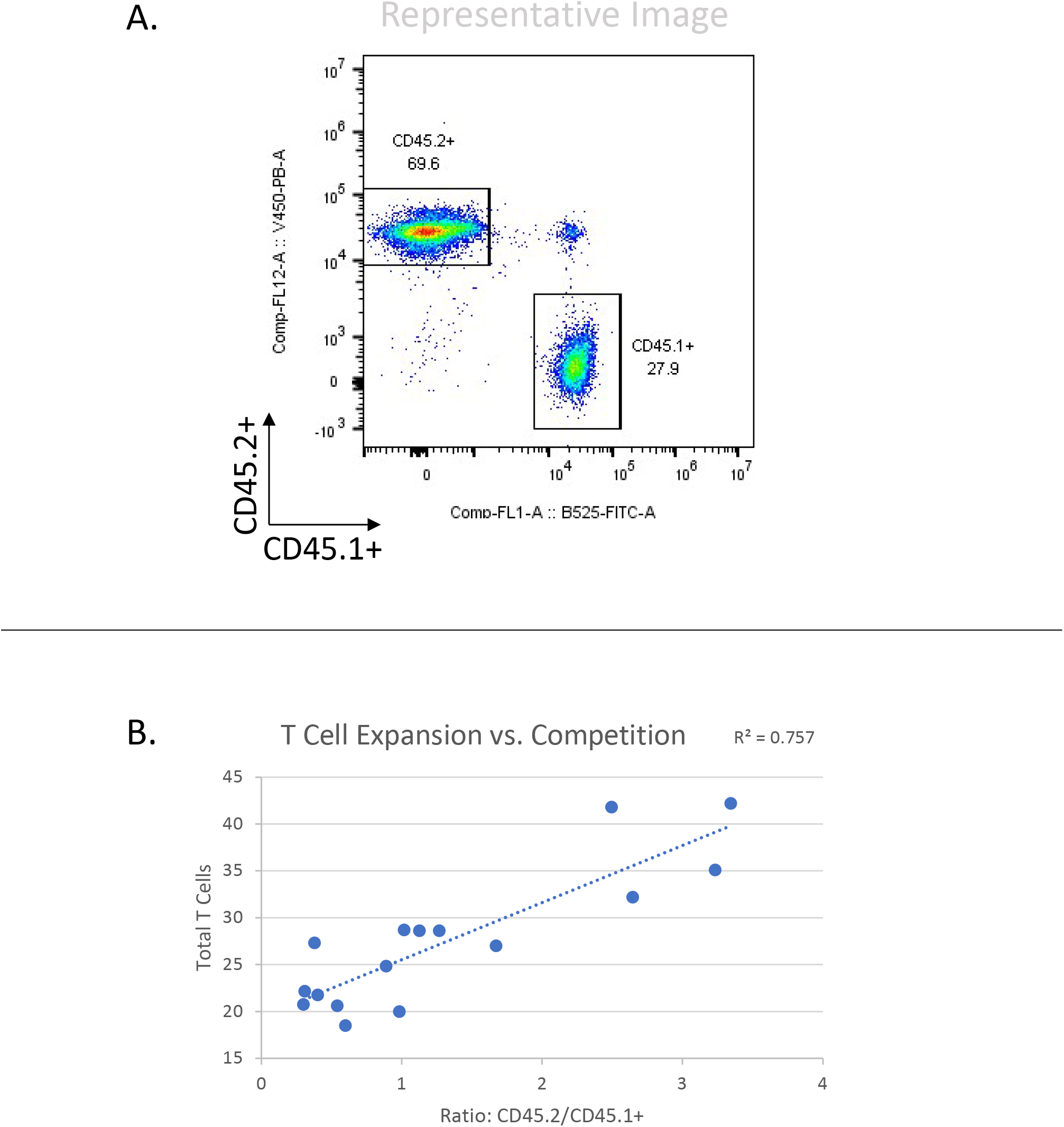
T cell expansion corresponds with expansion of p53-null cells. Positive correlation between *Trp53*^-/-^ leukocyte and T cell expansion across time points shown in Figure 1.

**Supplemental Figure 2:**
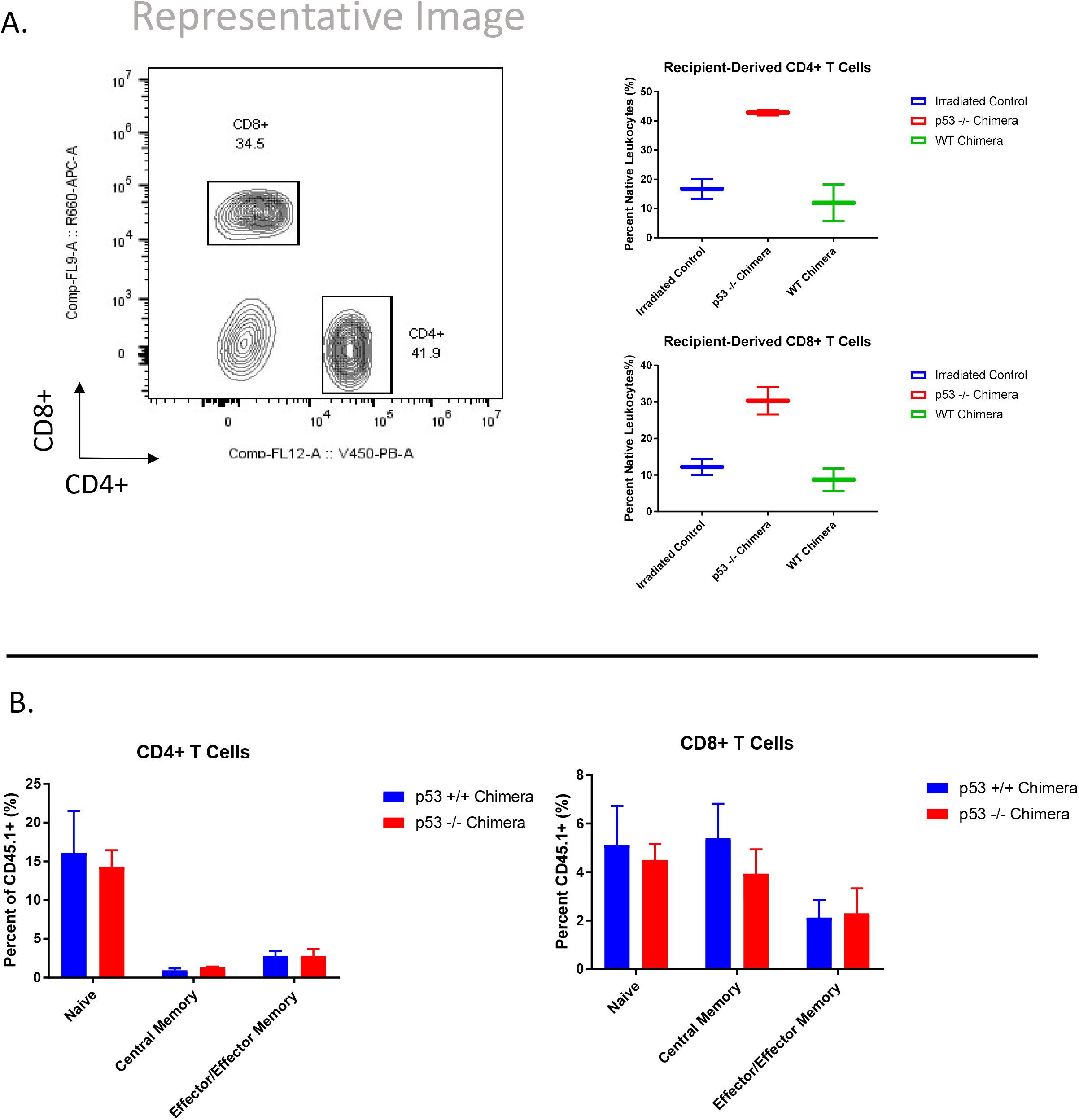
Expanded Recipient-Derived T Cells following sublethal Irradiation and Transplantation of KSL. (A) Expanded T cells at 12 weeks post-transplantation when recipients were sublethally irradiated and transplanted with cultured KSL. (B) CD4+ CD8+ T cell effector/effector memory differentiation at two weeks following transplantation with KSL.

**Supplemental Figure 3:**
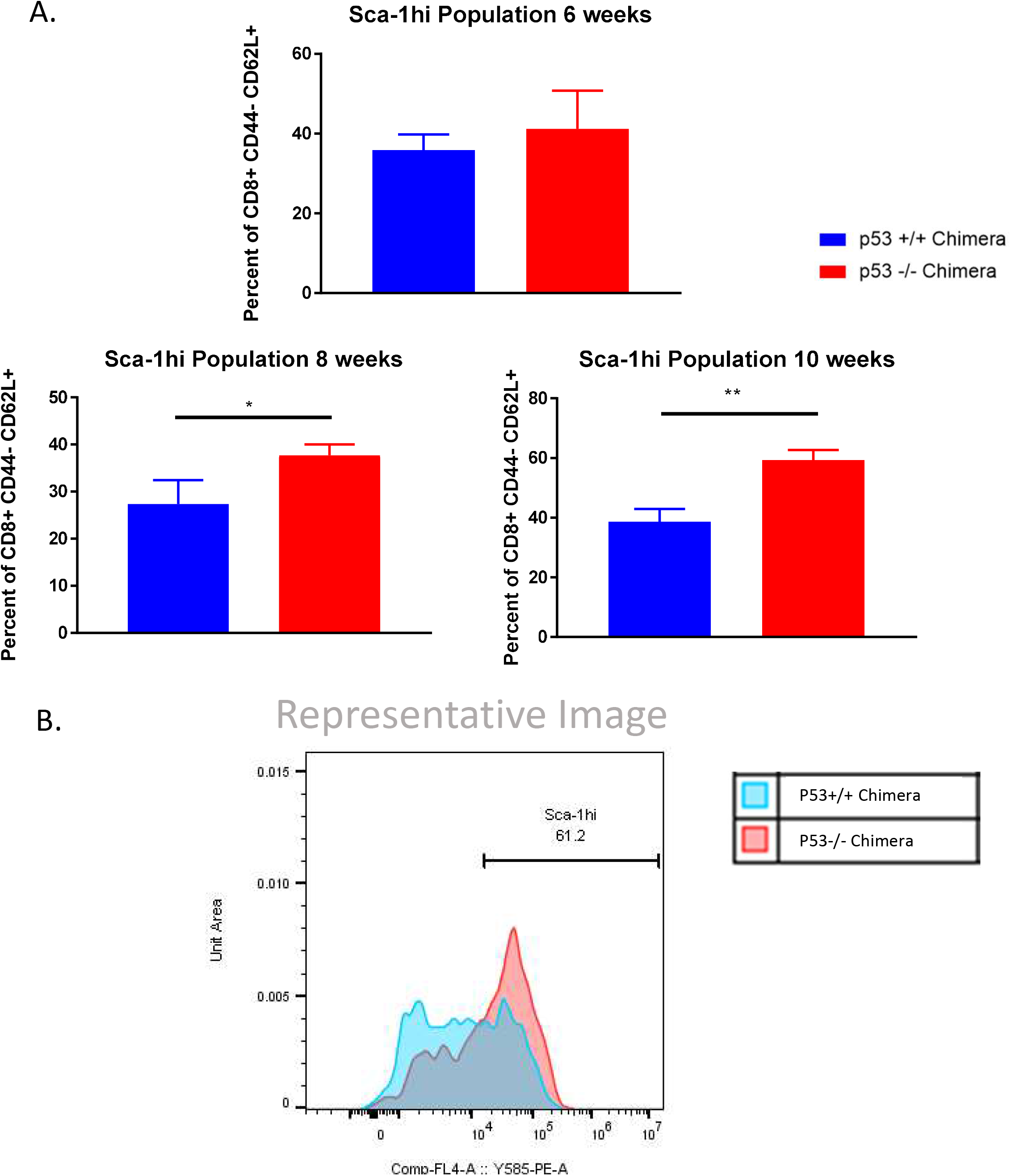
Upregulation of Sca-1 by CD8+ CD44-CD62L+ *Trp53* chimeras. (A) Expression of Sca-1 at time points indicated by sub-lethally irradiated mice transplanted with equivalent numbers of either *Trp53*^-/-^ or *Trp53*^+/+^ KSL. (B) Representative flow plot showing histograms

**Supplemental Figure 4:**
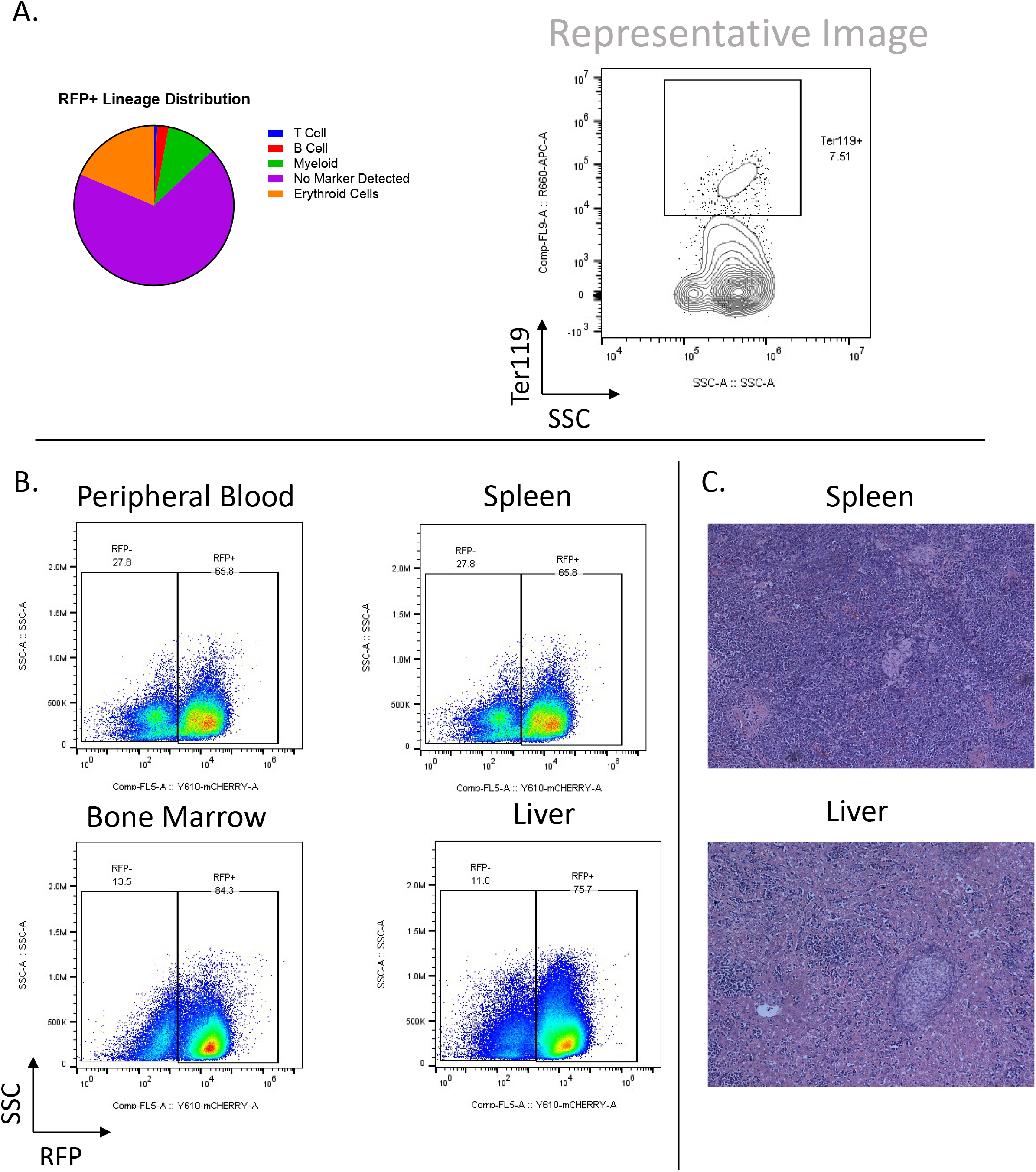
RFP+ Lineage Distribution with Erythroid compartment. (A) Immunophenotype of *Trp53*^-/-^;*Nras^G12D^* leukemia indicating minimal but present erythroid maturation. (B) Representative flow plots of various hematopoietic tissues and (C) corresponding H&E staining of spleen and liver

## References

1. Sperling, A.S., C.J. Gibson, and B.L. Ebert, The genetics of myelodysplastic syndrome: from clonal haematopoiesis to secondary leukaemia. Nature reviews. Cancer, 2017. 17(1): p. 5–19.

2. Chen, S. and Y. Liu, p53 involvement in clonal hematopoiesis of indeterminate potential. Curr Opin Hematol, 2019. 26(4): p. 235–240.

3. Bowen, D., et al., TP53 gene mutation is frequent in patients with acute myeloid leukemia and complex karyotype, and is associated with very poor prognosis. Leukemia, 2009. 23(1): p. 203–6.

4. Middeke, J.M., et al., TP53 mutation in patients with high-risk acute myeloid leukaemia treated with allogeneic haematopoietic stem cell transplantation. Br J Haematol, 2016. 172(6): p. 914–22.

5. Döhner, H., D.J. Weisdorf, and C.D. Bloomfield, Acute Myeloid Leukemia. 2015. 373(12): p. 1136–1152.

6. Hunter, A.M. and D.A. Sallman, Current status and new treatment approaches in TP53 mutated AML. Best Practice & Research Clinical Haematology, 2019. 32(2): p. 134–144.

7. Desai, P., et al., Somatic mutations precede acute myeloid leukemia years before diagnosis. Nat Med, 2018. 24(7): p. 1015–1023.

8. Wong, T.N., et al., Role of TP53 mutations in the origin and evolution of therapy-related acute myeloid leukaemia. Nature, 2015. 518(7540): p. 552–555.

9. Ok, C.Y., et al., TP53 mutation characteristics in therapy-related myelodysplastic syndromes and acute myeloid leukemia is similar to de novo diseases. Journal of hematology & oncology, 2015. 8: p. 45–45.

10. Jaiswal, S., et al., Clonal Hematopoiesis and Risk of Atherosclerotic Cardiovascular Disease. New England Journal of Medicine, 2017. 377(2): p. 111–121.

11. Sano, S., et al., Tet2-Mediated Clonal Hematopoiesis Accelerates Heart Failure Through a Mechanism Involving the IL-1β/NLRP3 Inflammasome. Journal of the American College of Cardiology, 2018. 71(8): p. 875–886.

12. Sano, S., et al., JAK2 (V617F) -Mediated Clonal Hematopoiesis Accelerates Pathological Remodeling in Murine Heart Failure. JACC. Basic to translational science, 2019. 4(6): p. 684–697.

13. Lim, G.B., Clonal haematopoiesis, aortic stenosis and reduced survival after TAVI. Nature Reviews Cardiology, 2019. 16(11): p. 650–650.

14. Rutella, S., et al. Immune infiltration correlates with TP53 mutational status in a multi-cohort acute myeloid leukemia study. in JOURNAL FOR IMMUNOTHERAPY OF CANCER. 2019. BMC CAMPUS, 4 CRINAN ST, LONDON N1 9XW, ENGLAND.

15. Danaher, P., et al., Gene expression markers of Tumor Infiltrating Leukocytes. J Immunother Cancer, 2017. 5: p. 18.

16. Tyner, J.W., et al., Functional genomic landscape of acute myeloid leukaemia. Nature, 2018. 562(7728): p. 526–531.

17. Ley, T.J., et al., Genomic and epigenomic landscapes of adult de novo acute myeloid leukemia. N Engl J Med, 2013. 368(22): p. 2059–74.

18. Zhang, J., et al., p53^-/-^ synergizes with enhanced NrasG12D signaling to transform megakaryocyte-erythroid progenitors in acute myeloid leukemia. Blood, 2017. 129(3): p. 358–370.

19. Wellenstein, M.D., et al., Loss of p53 triggers WNT-dependent systemic inflammation to drive breast cancer metastasis. Nature, 2019. 572(7770): p. 538–542.

20. Duan, J., Y. Wang, and S. Jiao, Checkpoint blockade-based immunotherapy in the context of tumor microenvironment: Opportunities and challenges. Cancer medicine, 2018. 7(9): p. 4517–4529.

21. Stahl, M. and A.D. Goldberg, Immune Checkpoint Inhibitors in Acute Myeloid Leukemia: Novel Combinations and Therapeutic Targets. Curr Oncol Rep, 2019. 21(4): p. 37.

22. Goltz, D., et al., PD-L1 (CD274) promoter methylation predicts survival in patients with acute myeloid leukemia. Leukemia, 2017. 31(3): p. 738–743.

23. Wang, X., et al., Tumor suppressor miR-34a targets PD-L1 and functions as a potential immunotherapeutic target in acute myeloid leukemia. Cell Signal, 2015. 27(3): p. 443–52.

24. Zajac, M., et al., Expression of CD274 (PD-L1) is associated with unfavourable recurrent mutations in AML. Br J Haematol, 2018. 183(5): p. 822–825.

25. Stubbs, M.C. and A.V. Krivtsov, Murine Retrovirally-Transduced Bone Marrow Engraftment Models of MLL-Fusion-Driven Acute Myelogenous Leukemias (AML). Current protocols in pharmacology, 2017. 78: p. 14.42.1–14.42.19.

